# Low spatial structure and selection against secreted virulence factors attenuates pathogenicity in *Pseudomonas aeruginosa*

**DOI:** 10.1101/250845

**Authors:** Elisa T. Granato, Christoph Ziegenhain, Rasmus L. Marvig, Rolf Kümmerli

**Affiliations:** Department of Plant and Microbial Biology, University of Zurich, Zurich, Switzerland; Department of Zoology, University of Oxford, Oxford, United Kingdom.; Department Biology II, Ludwig-Maximilians-University, Munich, Germany.; Department of Cell and Molecular Biology, Karolinska Institutet, Solna, Sweden.; Center for Genomic Medicine, Rigshospitalet, Copenhagen, Denmark.

## Abstract

Bacterial opportunistic pathogens are feared for their difficult-to-treat nosocomial infections and for causing morbidity in immunocompromised patients. Here, we study how such a versatile opportunist, *Pseudomonas aeruginosa*, adapts to conditions inside and outside its model host *Caenorhabditis elegans*, and use phenotypic and genotypic screens to identify the mechanistic basis of virulence evolution. We found that virulence significantly dropped in unstructured environments both in the presence and absence of the host, but remained unchanged in spatially structured environments. Reduction of virulence was either driven by a substantial decline in the production of siderophores (in treatments without hosts) or toxins and proteases (in treatments with hosts). Whole-genome sequencing of evolved clones revealed positive selection and parallel evolution across replicates, and showed an accumulation of mutations in regulator genes controlling virulence factor expression. Our study identifies the spatial structure of the non-host environment as a key driver of virulence evolution in an opportunistic pathogen.

## INTRODUCTION

Understanding how microbial pathogens evolve is essential to predict their epidemiological spread through host populations and the damage they can inflict on host individuals. Evolutionary theory offers a number of concepts aiming at forecasting the evolution of pathogen virulence and identifying the key factors driving virulence evolution [1,2]. While most evolutionary models agree that the spatial structure of the environment is an important determinant of virulence evolution, they differ on whether spatial structure should boost or curb pathogen virulence. One set of models predicts that high spatial structure lowers virulence, because it favors clonal infections and thereby limits the risk of hosts being infected by multiple competing pathogen lineages [3–6]. In this scenario, it is thought that the interests of pathogens in clonal infections become aligned, which should select for prudent host exploitation and thus low virulence [7,8]. Another set of models predicts that high spatial structure increases virulence because it favors the cooperative secretion of harmful virulence factors required for successful host colonization [5,9,10]. These models are based on the idea that virulence factors, such as toxins, proteases and iron-scavenging siderophores, are shared between pathogen individuals in infections [11–13]. Hence, low spatial structure is predicted to favor the evolution of cheating mutants that exploit the virulence factors produced by others, without contributing themselves [14]. Invasions of these cheats would then lower overall virulence factor availability and damage to the host [15–19].

Both classes of models have received some empirical support. While experimental evolution studies with viruses showed that limited dispersal indeed favors more benign pathogens [20–22], work with bacteria showed evidence for the opposite pattern [17,23,24]. Although these studies significantly advanced our understanding of virulence evolution, several fundamental questions remain still open. For instance, we generally know little about the mechanistic basis of virulence evolution [8,20,21,25]. Moreover, bacterial studies often built on controlled mixed versus monoinfections using wildtype strains and engineered mutants deficient for virulence factor production [17,23,24]. It thus remains unknown whether virulence-factor deficient mutants would indeed evolve *de novo* and spread to high frequency. Finally, we have limited understanding of how adaptation to the non-host environment affects virulence evolution [26,27], since most studies on bacterial opportunistic pathogens involved direct host-to-host transfers [28–30].

Here we aim to tackle these unaddressed issues by conducting an experimental evolution study, where we (i) allow opportunistic bacterial pathogens to adapt both to the host and the non-host environment, (ii) manipulate the spatial structure of the environment, and (iii) uncover the targets of selection and mechanisms provoking virulence change using high-throughput phenotypic screening combined with whole-genome sequencing of evolved clones. For our approach, we used the opportunistic human pathogen *Pseudomonas aeruginosa* infecting its model host, the nematode *Caenorhabditis elegans* [31,32]. This bacterium is typically acquired by the host from an environmental reservoir [33,34], and nematodes can quickly become infected through the intestinal tract because they naturally feed on bacteria [35]. In our experiment, we let *P. aeruginosa* PAO1 wildtype bacteria evolve for 60 days in four different environments in eight-fold replication, implementing a 2×2 full factorial design (Fig. 1A). To assess the role of spatial structure of the environment (first factor) for virulence evolution, we let the pathogens evolve in either unstructured uniform liquid or spatially structured solid medium. To understand how adaptation to the non-host environment affects virulence within the host, we further let the pathogens evolve both in the presence and the absence of the host (second factor). Following evolution, we quantified changes in pathogenicity for each independent replicate, and assessed whether these changes are associated with alterations in the expression of four important virulence factors of *P. aeruginosa*, which include the siderophore pyoverdine, the toxin pyocyanin, secreted proteases, and the ability to form biofilms. Finally, we whole-genome sequenced 140 evolved clones to map phenotypes to genotypes, and to test for positive selection, parallel evolution among independent replicates, and orders of mutations during evolution.

**Fig. 1.**
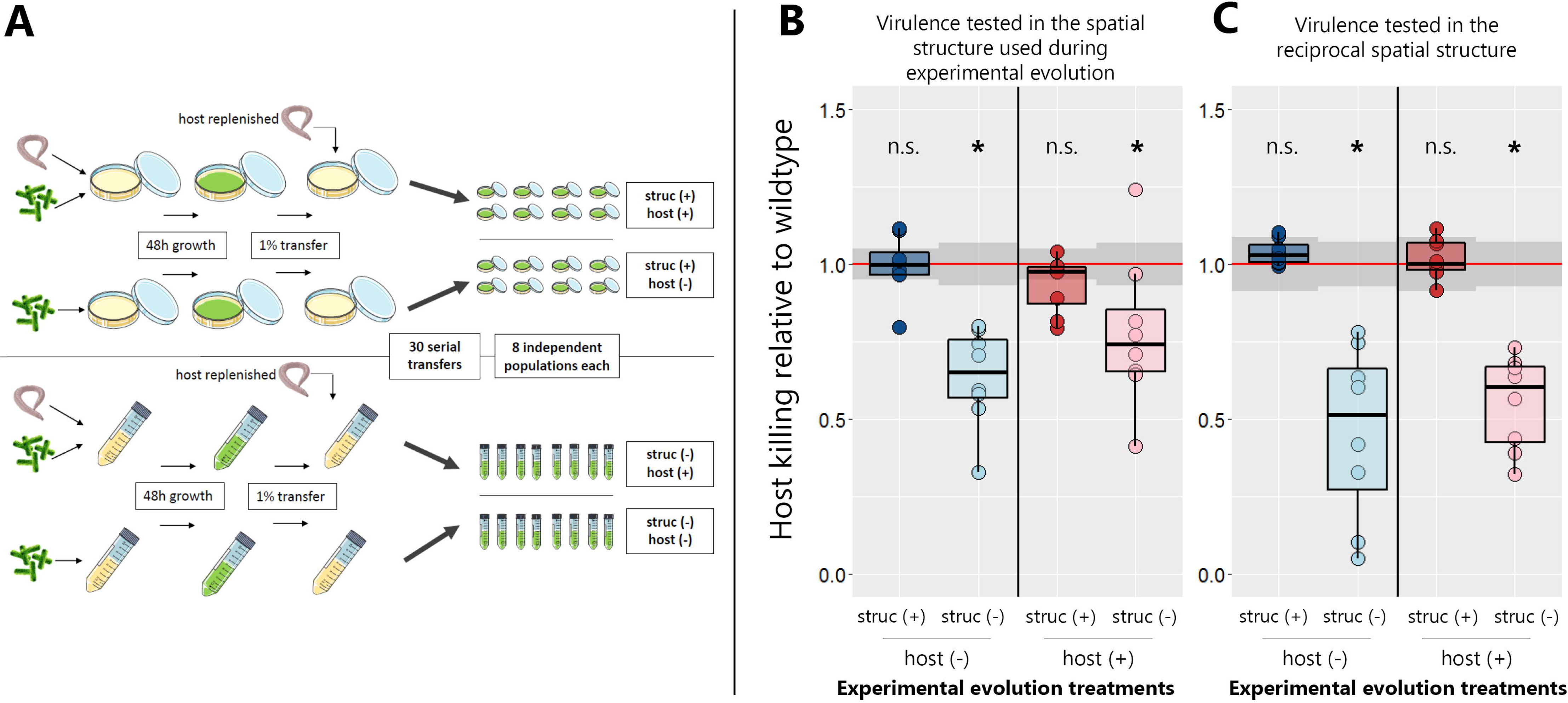
Virulence decreased during evolution in spatially unstructured environments. (**A**)Experimental design: *P. aeruginosa* PAO1 bacteria were serially transferred 30 times in four different environments in 8-fold replication. These environments were either spatially structured (“struc +”) or unstructured (“struc ‒”), and either contained (“host +”) or did not contain (“host ‒”) *C. elegans* nematodes for the bacteria to infect. Subsequently, the evolved populations were tested for their virulence towards the nematode under two different conditions: (**B**) In the environment the populations evolved in (i.e. populations that evolved on agar plates tested on agar plates, populations that evolved in liquid culture tested in liquid culture); and (**C**) in the reciprocal environment as a control (populations that evolved on agar plates tested in liquid culture, populations that evolved in liquid tested on agar plates). Both assays revealed that virulence significantly decreased during evolution in unstructured environments (Wilcoxon rank-sum test, asterisks denote p < 0.05; see Table S1). Virulence was quantified as percent nematodes killed at 24 h post infection, scaled to the ancestral wildtype. Individual dots represent mean virulence of evolved populations across three replicates. The red line represents the average wildtype virulence level in the respective assay, with shaded areas denoting the 95% confidence intervals.

## MATERIALS AND METHODS

### Strains and culturing conditions

*Pseudomonas aeruginosa* wildtype strain PAO1 (ATCC 15692) constitutively expressing GFP (PAO1-*gfp*) was used for experimental evolution. The siderophore-deficient mutant PAO1Δ*pvdD-gfp*, the quorum-sensing deficient mutants PAO1Δ*rhlR* and PAO1Δ*lasR* (S. Diggle, Georgia Institute of Technology, USA), and the biofilm-deficient mutant MPAO1Δ*pelA*Δ*pslA* (M. Toyofuku, University of Zurich, Switzerland) were used as negative controls for phenotype screening. For overnight pre-culturing, we used Lysogeny Broth (LB) and incubated cultures under shaking conditions (190 rpm) for 18-20 h. Optical density (OD) of bacterial cultures was determined in a Tecan Infinite M-200 plate reader (Tecan Group Ltd., Switzerland) at a wavelength of 600 nm. All experiments were conducted at 25°C, except for pre-culturing of the ancestral strain before experimental evolution (see below). To generate iron-limited nutrient medium (RDM-Ch) suitable for bacterial and nematode co-culturing, we supplied low-phosphate NGM (nematode growth medium; 2.5 gL^−1^ BactoPeptone, 3 gL^−1^ NaCl, 5 mgL^−1^ Cholesterol, 25 mM MES buffer pH = 6.0, 1mM MgSO4, 1mM CaCl_2_; adapted from [32]) with 200 μM of the iron chelator 2,2’-Bipyridyl. For agar plates, media were supplemented with 1.5% (m/V) agar. All chemicals were acquired from Sigma-Aldrich, Switzerland. *Caenorhabditi elegans* N2 wildtype nematodes were acquired from the *Caenorhabditis Genetics Center.* Nematode maintenance and generation of age-synchronized nematodes was performed according to standard protocols [36].

### Experimental evolution

Experimental evolution was started with a clonal population of PAO1-*gfp*. For each of the four experimental treatments (agar plates with and without host, liquid culture with and without host), eight replicate lines were evolved independently (Fig. 1). During experimental evolution, *C. elegans* was not allowed to co-evolve. Instead, fresh L4-stage nematodes were supplied at each transfer step. Since *P. aeruginosa* is highly virulent towards *C. elegans*, the vast majority of worms were dead before each transfer step and we never observed any live larvae.

To start the experimental evolution, pre-cultures of PAO1-*gfp* were washed, OD-adjusted and either spread onto RDM-Ch agar plates or inoculated into liquid RDM-Ch in culture tubes (Fig. 1). For the “with host” treatments, L4-stage *C. elegans* nematodes were then added to each plate/culture tube (details in Supplementary Material). Plates and culture tubes were incubated for 48 h before the first transfer. Transfers of bacteria to fresh nutrient medium and, if applicable, addition of fresh nematodes to the samples were conducted every 48 h (details in Supplementary Material). Briefly: bacteria were transferred by replica-plating (for “agar plate” treatments) and a fraction of nematodes was carried over (for “agar plates with host” treatments); or bacteria were transferred by inoculating fresh media with an aliquot from the previous culture (for “liquid” treatments), and a fraction of nematodes was carried over (for “liquid with host” treatments). The number of viable bacteria transferred through replica-plating corresponded approximately to a 1:100 dilution, and was therefore equivalent to the dilution achieved in the liquid cultures. In total, 30 transfers were conducted, corresponding to approximately 200 generations of bacterial evolution. At the end of the experimental evolution, evolved populations were frozen at −80°C.

### Killing assays

Population level virulence was assessed in two different killing assays at 25°C, namely in liquid culture and on agar plates, representing the two different environments the bacterial populations evolved in. Populations were separately tested both in the environment they evolved in (populations evolved on agar plates tested on agar plates, and populations evolved in liquid culture tested in liquid culture), and in the respective reciprocal environment (populations evolved in liquid culture tested on agar plates, and vice versa).

For killing assays in liquid culture, evolved bacterial populations and the ancestral wildtype were inoculated into liquid RDM-Ch in three replicate culture tubes per population. After an incubation period of 48 h, ~2500 L4-stage nematodes were added, and culture tubes further incubated for 48 h. Virulence was determined by counting the fraction of dead worms at 24 h and 48 h following nematode addition. For killing assays on agar plates, evolved bacterial populations and the ancestral wildtype were spread on six replicate RDM-Ch agar plates per population. Plates were then incubated for 48 h, and 20-60 L4-stage nematodes were added to the plates. Virulence was determined by counting the fraction of dead worms at 24 h and 48 h after adding the nematodes. More details on the killing assays can be found in the Supplementary Material.

### Phenotypic screening of single clones

Evolved bacterial populations were re-grown from freezer stocks and twenty colonies were randomly isolated for each population. In total, 640 clones were isolated and subjected to phenotypic screens for virulence factor production. Pyoverdine production was measured in liquid RDM-Ch in 96-well plates. Plates were incubated for 24 h under shaken conditions and OD600 and pyoverdine-specific fluorescence (ex: 400 nm / em: 460 nm) were measured in a plate reader. Pyocyanin production was measured in liquid LB in 24-well plates. Plates were incubated for 24 h under shaken conditions, and pyocyanin was quantified by measuring OD at 691 nm of the cell-free supernatant in a plate reader. Protease production was measured using skim milk agar in 24-well plates. 1 μL of bacterial culture was dropped onto the agar, and plates were incubated for 20 h. Protease production was quantified by measuring the resulting halo with ImageJ [37]. Biofilm production was measured in liquid LB in 96-well plates. Plates were incubated under static conditions for 24 h, and the production of surface-attached biofilms was quantified by calculating the “Biofilm Index” (OD570/OD550) for each well [38]. Details on the phenotypic screening can be found in the Supplementary Material.

### Calculation of the “virulence factor index”

We defined a virulence factor index *v* = Σ *r*_i_ / *n*, where *r*_i_-values represent the average virulence factor production scaled relative to the ancestral wildtype for the *i*-th virulence factor (*i* = pyoverdine, pyocyanin, proteases, biofilm), and *n* is the total number of virulence factors. For clones with wildtype production levels for all four virulence factors, *v* = 1, whereas *v* < 1 would represent clones with overall reduced production levels. For statistical analyses and data presentation, we used the average virulence index across clones for each population.

### Whole-genome sequencing of evolved clones

To select populations and clones for sequencing, we first chose all populations with decreased virulence, and then added randomly chosen populations to cover all four treatments in a balanced way (four sequenced populations per treatment), leading to a total of 16 selected populations. From these, we selected nine clones per population according to the following scheme: first, we tried to get at least one clone that showed no phenotypic differences to the ancestral wildtype with regards to pyoverdine and pyocyanin production. Then, we tried to get clones with a marked decrease in pyoverdine and/or pyocyanin production. Finally, we filled up the list with randomly chosen clones. Genomic DNA was isolated from all selected clones using a commercial kit, sequencing libraries were constructed using the Nextera XT Kit (Illumina, USA) and whole-genome sequencing was performed 2×150 bp on a NextSeq500 (Illumina, USA; details in Supplementary Material).

### Variant calling

Demultiplexed reads were aligned to the *P. aeruginosa* PAO1 reference genome using bowtie2 in local-sensitive mode [39]. PCR duplicates were removed using “picard” tools (https://broadinstitute.github.io/picard/). Variants were called using “samtools” (v0.1.19), “mpileup” and “bcftools” [40] and filtered with default parameters using “samtools” and “vcfutils”. Variant effects were predicted using SnpEff (version 4.1d) [41]. Detailed protocols for variant analysis and phylogenetic inference are provided in the Supplementary Material.

### Statistical Analysis

We used linear models and linear mixed models for statistical analyses in R 3.2.2 [42]. When data distributions did not meet the assumptions of linear models, we performed non-parametric Wilcoxon rank-sum tests. To test whether virulence factor production in single clones depended on the environment they evolved in, we used Markov-chain Monte Carlo generalized linear mixed models (MCMCglmm) [43]. Principal component analysis (PCA) was conducted using the ‘FactoMineR’ [44] and ‘factoextra’ packages (https://CRAN.R-project.org/package=factoextra). Detailed description of statistical methods and test results are provided in the Supplementary Methods and Table S1.

## RESULTS

### Selection for reduced virulence in environments with low spatial structure

Prior to experimental evolution, we found that the ancestral wildtype was highly virulent by killing 76.2% and 83.9% of all host individuals within 24 hours in liquid and on solid media, respectively (Table S2). This pattern changed during evolution in spatially unstructured environments, where virulence dropped by 32.3% and 44.7% for populations that evolved with and without hosts, respectively (Fig. 1B+C, Fig. S1). Conversely, virulence remained high in structured environments. Overall, there was a significant effect of spatial structure on virulence evolution (linear mixed model: df_structure_ = 24.7, *t*_structure_ = - 2.11, *p*_structure_ = 0.045), while host presence did not seem to matter (df_host_ = 18.6, *t*_host_ = 0.86, *p*_host_ = 0.40).

### Treatment-specific changes in virulence factor production

To explore whether shareable virulence factors were under selection and whether changes in virulence factor production could explain the evolution of virulence, we isolated 640 evolved clones and quantified their production of: (i) pyoverdine, required for iron-scavenging [45]; (ii) pyocyanin, a broad-spectrum toxin [46]; and (iii) proteases to digest extracellular proteins [47]. We further quantified the pathogens’ ability to form biofilms on surfaces, another social trait typically involved with virulence [48]. We focussed on these four virulence-related traits because of their demonstrated relevance in the *C. elegans* infection model [32,48–50].

Our phenotype screens revealed significant treatment-specific changes in the production of all four virulence factors (Fig. 2). For pyoverdine, we observed that production levels of evolved clones were significantly lower in the unstructured environments without hosts compared to the other treatments (Fig. 2A; Bayesian generalized linear mixed model, BGLMM, significant interaction: *p*_host:structure_ = 0.027). Production levels were lower because many clones (44.4%) have partially or completely lost the ability to produce pyoverdine (Fig. 2A). Since these mutants appeared in six out of eight replicates (Fig. S2) and our media was iron-limited, impeding the growth of pyoverdine non-producers, these clones likely represent social cheaters, exploiting the pyoverdine secreted by producers [51,52]. While mutants with abolished pyoverdine production also emerged in the unstructured environment with hosts, their frequency was much lower (5.0%).

**Fig. 2.**
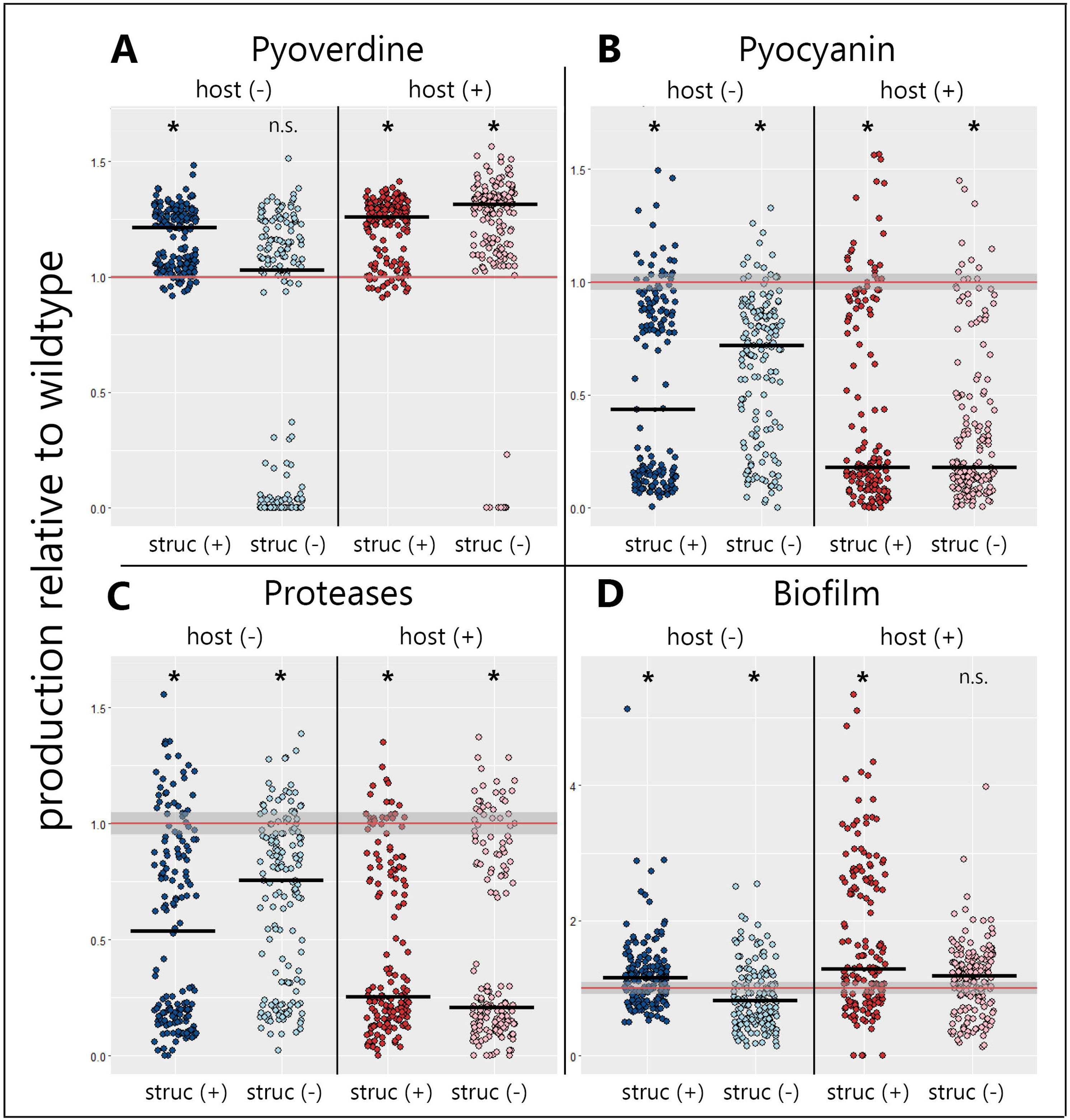
Selection promoted shifts in virulence factor production during experimental evolution. The production levels of four important virulence factors were determined for 640 evolved *P. aeruginosa* clones (20 clones per evolved line), and compared to the ancestral wildtype (mean ± 95 % confidence intervals indicated as red lines and shaded areas, respectively). (**A**) Pyoverdine production significantly increased in all treatments, except in the host-free unstructured environment, where 44% of all evolved clones partially or completely lost the ability to produce this siderophore. (**B**) The production of the toxin pyocyanin significantly decreased in all environments, but more so in the environments with the host. (**C**) The production of proteases also significantly decreased in all environments, with a sharper decline in environments with the host. (**D**) The clones’ ability to form surface-attached biofilms significantly decreased in the unstructured host-free environment, but significantly increased in all other environments. host (‒) = host was absent during evolution; host (+) = host was present during evolution; struc (‒) = evolution in a liquid-shaken unstructured environment; struc (+) = evolution in a structured environment on agar. We used non-parametric Wilcoxon rank-sum test for comparisons relative to the ancestral wildtype (asterisks denote p < 0.05), and BGLMM to test for treatment effects (see Table S1). Solid black bars denote the median for each treatment.

Pyocyanin production, meanwhile, significantly dropped in all four environments (Fig. 2B), but more so in the presence than in the absence of the host (*p*_host_ = 0.038), while spatial structure had no effect (*p*_structure_ = 0.981). The pattern of evolved protease production mirrored the one for pyocyanin (Fig. 2C): there was a significant overall decrease in protease production, with a significant host (*p*_host_ = 0.042), but no structure (*p*_structure_ = 0.489) effect. Since neither pyocyanin nor proteases are necessary for growth in our media, consisting of a protein-digest, reduced expression could reflect selection against dispensable traits. During infections, however, these traits are known to be beneficial [49,50] and accelerated loss could thus be explained by cheating, as secreted virulence factors could become exploitable inside the host. It is known that protease production can be exploited by non-producing clones [47], and there is recent evidence that the same might apply to pyocyanin [53]. The strong correlation between the pyocyanin and protease phenotypic patterns is perhaps not surprising, given that they are both regulated by the hierarchical quorum-sensing system of *P. aeruginosa* [54].

Finally, the clones’ ability to form surface-attached biofilms significantly increased in the presence of the host (*p*_host_ = 0.007) and in structured environments (*p*_structure_ = 0.010; Fig. 2D). These findings indicate that attachment ability might be less important under shaken conditions, but relevant within the host to increase residence time.

### Aggregate change in virulence factor production correlates with evolved virulence

While the phenotypic screens revealed altered virulence factor production levels, with significant host and environmental effects (Fig. 2), the virulence data suggest that there is no host effect, and spatial structure is the only determinant of virulence evolution (Fig 1). In the attempt to reconcile these apparently conflicting results, we first performed a principal component analysis (PCA) on population averages of the four virulence factor phenotypes (Fig. 3A). The PCA indicates that each treatment evolved in a different direction in phenotype space, a pattern confirmed by a PERMANOVA statistical analysis testing for spatial separation of treatment groups (*p* = 0.002). From this, we can deduct that environmental and host factors indeed both seem to matter. This analysis also shows that the direction of phenotypic changes was aligned for some traits, but opposed for others (Fig. 3A, Fig. S3). A decrease in pyocyanin production was generally connected to a decrease in protease production (Fig. S3D). On the other hand, decreased pyocyanin and protease production were associated with both higher pyoverdine and biofilm production (Fig. S3B+E).

**Fig. 3.**
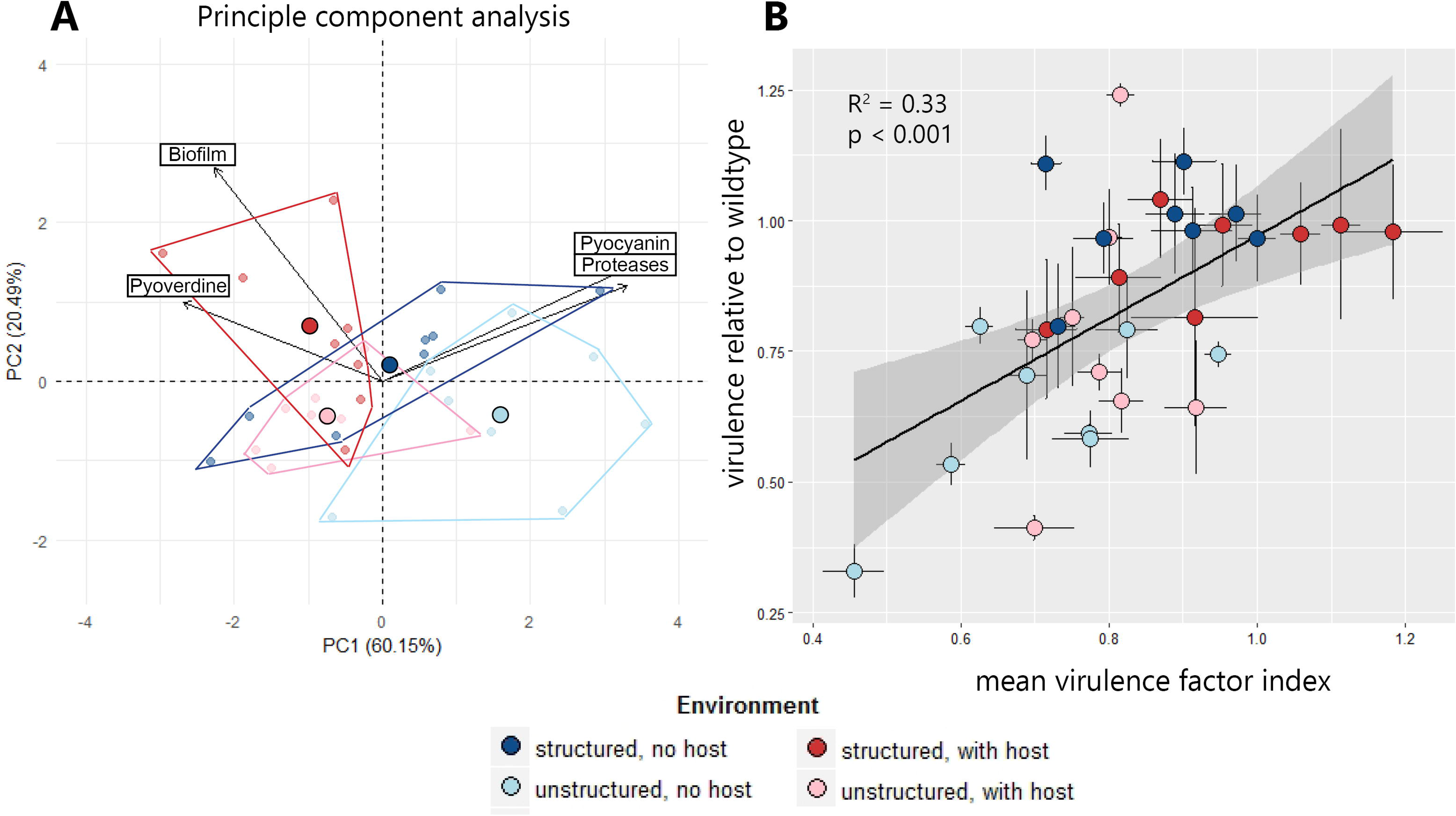
The aggregate change in virulence factor production explains virulence evolution. (**A**) A principal component analysis (PCA) on the population-level changes in the production of four virulence factors (pyoverdine, pyocyanin, proteases, biofilm) reveals divergent evolutionary patterns. For instance, analysis of the first two principal components (explaining 80.6 % of the total variance) shows complete segregation between populations evolved in unstructured host-free environments and structured environments with the host. Moreover, the PCA reveals that evolutionary change was aligned for some traits (aligned vectors for pyocyanin and proteases), but opposed for others (inversed vectors for pyoverdine versus pyocyanin/proteases). Small and large symbols depict individual population values and average values per environment, respectively. Polygons show the boundaries in phenotype space for each environment. (**B**) We found that the aggregate change in the production of all four virulence factors explained the evolution of virulence. To account for the aligned and opposing effects revealed by the PCA, we defined the “virulence factor index” as the average change in virulence factor production across all four traits, scaled relative to the ancestral wildtype. Symbols and error bars depict mean values per population and standard errors of the mean, respectively.

Given these opposing evolutionary directions and trade-offs between virulence factors we hypothesized that an increase in the production of one virulence factor could (at least partially) be counterbalanced by the reduction of another virulence factor. In the extreme case, two virulence factors could both be under selection, but in opposite directions, such that their net effects on virulence could cancel out. In line with this hypothesis, we found that the evolutionary change in virulence could only be explained when considering the aggregate change of all virulence factor phenotypes (Fig. 3B, R^2^ = 0.33, F(1,30) = 14.7, *p* < 0.001; also see Fig. S4), but not when focussing on single virulence factors (Fig. S5). Thus, decreased virulence in unstructured environments is attributable to a simultaneous decrease in the production of multiple virulence factors (i.e. pyocyanin, proteases, and sometimes pyoverdine). Conversely, unchanged virulence in structured environments can be explained by compensatory effects (i.e. the reduction in pyocyanin and protease production is balanced by increased pyoverdine and biofilm production). Important to note is that the observed pyoverdine upregulation is presumably a compensatory phenotypic response, as decreased pyocyanin and protease production are known to lower iron availability [55], which in turn might trigger increased pyoverdine production.

### Mutations in key regulators explain changes in virulence factor phenotypes

To examine whether genetic changes can explain the observed shifts in virulence factor production, we successfully sequenced the genome of 140 evolved clones from 16 independent populations and compared them to the ancestor. Relative to the ancestral wildtype, we identified 182 mutations (153 SNPs and 29 microindels, i.e. small insertions and deletions), with 5-49 mutations per population (median = 8.5). Individual clones accumulated 0-5 mutations, except for one clone (PA-030) with 42 mutations, of which 41 mutations were in a 5022 bp Pf1 prophage region, a known mutational hotspot [56].

We identified 18 loci (genes and intergenic regions) that were independently mutated in at least two populations (Fig. 4A). The most frequently mutated gene was *lasR*, encoding the regulator of the Las quorum-sensing (QS) system. The second most frequent mutational target were ten different *pil* genes, involved in type IV pili biosynthesis and twitching motility. The frequent mutations in this cluster suggest that mutations in any of these genes could potentially lead to a similar beneficial phenotype. Finally, the *pvdS* coding region or the *pvdG-pvdS* intergenic region, containing the *pvdS* promoter, were also often mutated (i.e. in five populations). PvdS is the iron-starvation sigma factor controlling pyoverdine synthesis, and mutations in this gene can lead to pyoverdine deficiency [19,52].

**Fig. 4.**
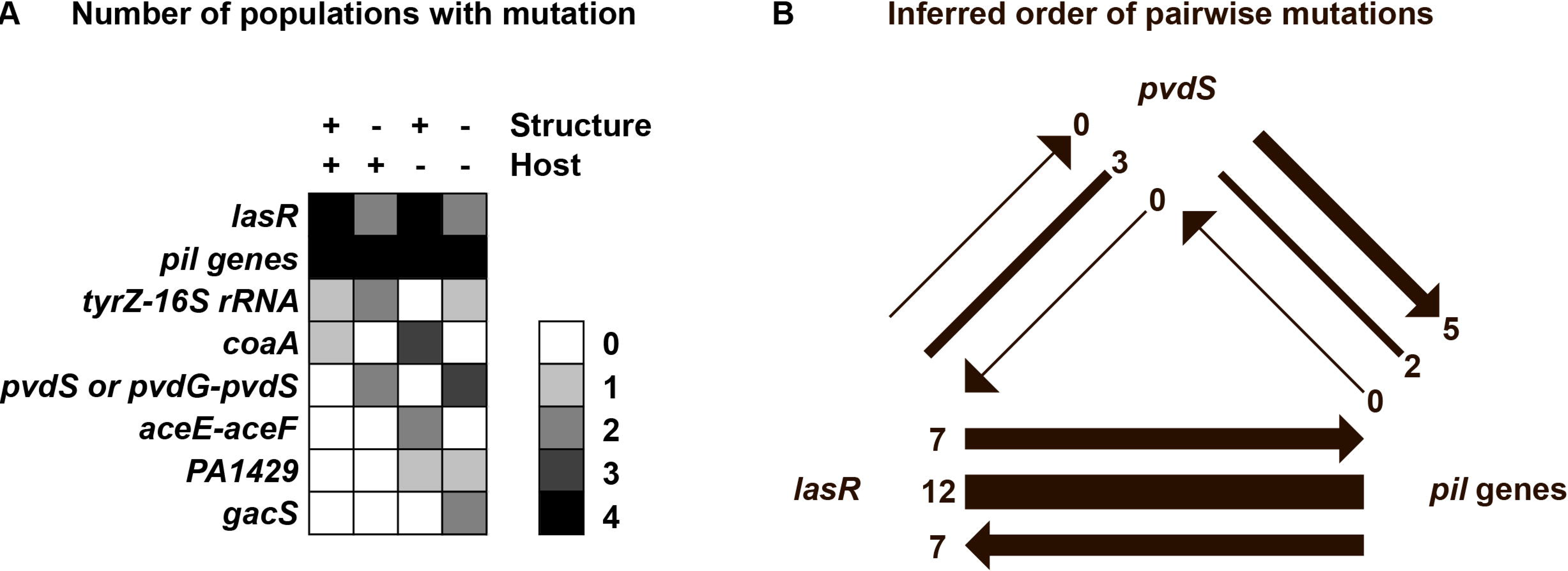
Whole genome sequencing reveals mutational profiles and order of mutations. Whole genomes of 140 evolved clones (four populations per environment and eight to nine clones per population) were sequenced, and SNPs and INDELs in genes and intergenic regions were called relative to the ancestral wildtype. **(A)** List of the loci that harbored mutations in at least two populations. The scale of grey shadings corresponds to the number of populations from each experimental condition in which clones with mutations in the respective loci occurred. (**B**) Phylogenetic interference of the order of mutations among clones harboring mutations in two of the most frequently affected loci. Order of mutations are indicated by arrows pointing towards the loci that were mutated second. Lines without arrowheads indicate that phylogenetic inference could not resolve the order of mutations.

We found that two of these frequently mutated targets explained a large proportion of the altered virulence factor phenotypes (Fig. 5). Specifically, reduced pyoverdine production was significantly associated with mutations in the *pvdS* gene or its promoter region (F(1,137) = 240.1, *p* < 0.0001, Fig. 5A). Moreover, there were significant correlations between reduced pyocyanin and protease production and mutations in *lasR* (pyocyanin: F(1,137) = 18.76, *p* < 0.0001; proteases: F(1,137) = 16.04, *p* < 0.001, Fig. 5B+C). In roughly half of the clones (pyocyanin: 51.3%, proteases: 45.6%), reduced production levels could be attributed to mutations in *lasR*. While the Las-system directly controls the expression of proteases, pyocyanin is only indirectly linked to this QS-system, via the two subordinate Rhl and PQS quorum sensing systems [54]. We further analyzed whether the mutations in the type IV pili genes affected biofilm formation. Although type IV pili can be important for bacterial attachment to surfaces [57], there was no clear relationship between these mutations and the evolved biofilm phenotypes (Fig. S6). This is probably because biofilm formation is a quantitative trait, involving many genes, and because we found both evolution of increased and decreased biofilm production, which complicates the phenotype-genotype matching. Alternatively, it could also be that the observed mutations rather affect twitching motility than biofilm formation, a trait we did not examine here, but might also be involved in virulence [58].

**Fig. 5.**
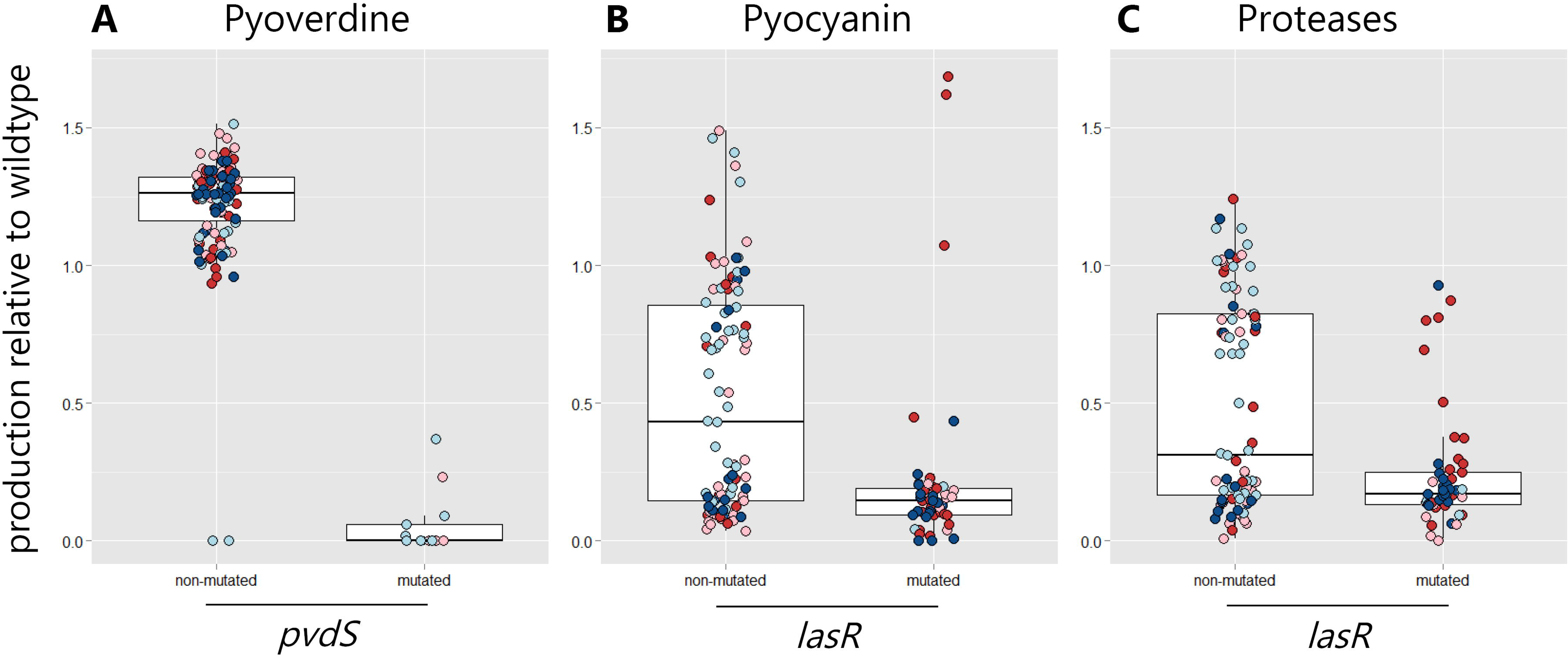
Mutations in key regulatory genes underlie the loss of virulence factor production. Across the 140 sequenced clones, there was an accumulation of mutations in two regulatory genes (*pvdS* and *lasR*), which significantly correlated with the phenotypic changes observed for pyoverdine (**A**), pyocyanin (**B**) and protease (**C**) production. *pvdS* encodes the iron starvation sigma factor and all clones with mutations in this gene or its promoter showed significantly impaired pyoverdine production. *lasR* encodes the regulator of the Las-quorum-sensing system, which directly controls the expression of several proteases. All clones with *lasR* mutations showed reduced protease production. The LasR regulator also has downstream effects on the Rhl- and PQS quorum-sensing systems, which control pyocyanin production. Consistent with this view, most clones with *lasR* mutations (93.8 %) showed decreased pyocyanin production. Although the genotype-phenotype match was nearly perfect for mutated clones, a considerable amount of clones also showed altered phenotypes without mutations in these two regulators, suggesting that some of the phenotypic changes are caused by mutations in yet unidentified loci.

### Mutational patterns reveal evidence for positive selection and parallel evolution

To test whether the mutated loci were under positive selection, we calculated the relative rates of nonsynonymous to synonymous SNPs (dN/dS) for loci mutated in at least two populations and for loci mutated only once. We found dN/dS = 6.2 for loci mutated in parallel in multiple populations, suggesting significant positive selection (P(X≥74)~pois(λ=12) < 0.0001, where λ is the expected number of nonsynonymous SNPs under neutral evolution and X is the observed number of nonsynonymous SNPs). Conversely, dN/dS = 0.3 for loci mutated in only a single population, indicating that these loci were under negative selection (P(X≤26)~pois(λ=87) < 0.0001). Altogether, our findings reveal that the 18 loci with multiple mutations underwent adaptive parallel evolution.

Finally, we used phylogenetic inference to resolve the order of mutations involving the *lasR*, *pvdS*, and *pil* genes (Fig. 4B, Table S3). Such analyses could reveal whether selection of mutations in certain genes is dependent on previous mutations in other genes. When analyzing evolved clones that mutated in at least two of these loci, we observed no clear patterns of dependencies in the order of mutations in *lasR-pil*-mutants and *lasR-pvdS*-mutants. For *pvdS-pil*-mutants, meanwhile, we found that mutations in *pvdS* tended to precede the mutations in *pil* genes. While sample size is too low to draw any strong conclusions, this observation could indicate that mutations in type IV pili are particularly beneficial in a pyoverdine-negative background.

## DISCUSSION

Using the opportunistic human pathogen *P. aeruginosa*, we show that bacterial virulence can evolve rapidly during experimental evolution, as a result of adaptation to both the host and the non-host environment. Overall, we found that *P. aeruginosa* evolved greatly reduced virulence in liquid unstructured environments, but remained highly virulent in spatially structured environments, regardless of whether its nematode host was present or absent. Phenotypic and genotypic screens provide strong evidence for positive selection on bacterial virulence factors and parallel adaptive evolution across independent replicates. Virulence reduction in unstructured environments without hosts was driven by a sharp decline in the production of the siderophore pyoverdine, and moderate decreases in protease and pyocyanin production. Conversely, virulence reduction in unstructured environment with hosts is explained by a stark decrease in protease and pyocyanin production, but not pyoverdine. Although the traits under selection seem to vary as a function of host presence, our findings are in strong support of evolutionary theory predicting that low spatial structure should select for reduced pathogenicity if virulence is mediated by secreted compounds such as toxins, proteases or siderophores [5,9]. The reason for this is that secreted virulence factors can be shared between cells, and can thus become exploitable by cheating mutants that no longer contribute to costly virulence factor production, yet still capitalize on those produced by others. The spread of such mutants is predicted to reduce overall virulence factor availability and to curb virulence, exactly as observed in our study.

Our results highlight how an in-depth mechanistic analysis of the traits under selection can deepen our understanding of virulence evolution. In the absence of our phenotypic and genetic trait analysis, we would be tempted to conclude that the presence of the host has no effect on virulence evolution, and that evolutionary change is entirely driven by the external non-host environment (Fig. 2). Our mechanistic trait analysis shows that such conclusions would be premature and yields several novel nuances of virulence evolution. First, we observed strong selection for pyoverdine-negative mutants only in the absence but not in the presence of the host (Fig. 2A). Pervasive selection against pyoverdine in unstructured, yet iron-limited medium, has previously been attributed to cheating [14]. Here, we show that the spread of pyoverdine nonproducers is apparently prevented in the presence of the host. One reason for this host-specific effect might be that the spatial structure inside hosts counteracts the selective advantage nonproducers experience outside the host. Second, we found that the presence of the host had a significant effect on the strength of selection against pyocyanin and protease production (Fig. 2B+C). We speculate that the presence of the host alters the reason for why these two virulence factors are selected against. In the absence of the host, neither pyocyanin nor proteases are required for growth, and their decline could be explained by selection against superfluous traits. Conversely, these two traits become beneficial in the presence of the host [49,50], such that selection against them could at least partially be explained by cheating. Third, we found evidence that the presence of the host selected for mutants with increased capacities to form biofilms (Fig. 2D). Apart from increasing residency time within hosts, the shift from a planktonic to a more sessile lifestyle typically goes along with fundamental changes in gene expression patterns [59,60], which might in turn affect virulence. Finally, we found that virulence factors were also under selection in treatments where the overall virulence level did not change (i.e. in structured environments). In these environments, however, reduced production of one virulence factor (e.g. protease and pyocyanin) was often compensated by the upregulation of other virulence factors (e.g. pyoverdine and biofilm), resulting in a zero net change in virulence.

A number of previous studies showed that when competition between virulence-factor producing and engineered non-producing bacteria is allowed for, then non-producing strains can often invade pathogen populations and thereby lower virulence [15,17,23,24]. While our work is in line with these findings, it makes several additional contributions. First, our experiment started with fully virulent clonal wildtype bacteria, and any virulence-factor deficient mutants had to evolve from random mutations and invade pathogen populations from extreme rarity. Hence, our study proves that the predicted mutants indeed arise *de novo* and are promoted by natural selection in independent parallel replicates. Second, our results highlight that multiple social traits are under selection simultaneously, which can lead to either additive effects (when traits are regulatorily linked, e.g. proteases and pyocyanin) or compensatory effects (when traits evolve in opposite directions, e.g. increased biofilm versus decreased protease production). Third, our study design captured the cycling of an opportunistic pathogen through the host and the non-host environment, as it would occur under natural conditions [26,27], an approach that allowed us to discover accidental virulence effects that are purely driven by adaptation to the non-host environment.

At the genetic level, our findings closely relate to previous work that has identified *lasR* as a key target of evolution in the context of chronic *P. aeruginosa* infections in the cystic fibrosis lung [61–65], in non-cystic fibrosis bronchiectasis [66], as well as in acute infections [18,29]. While the ubiquitous appearance of *lasR*-mutants was often interpreted as a specific host adaptation, we show here that *lasR*-mutants frequently arise even in the absence of a host, indicating that mutations in *lasR* are not a host-specific phenomenon. We propose three mutually non-exclusive explanations for the frequent occurrence and selective spread of *lasR*-mutants. First, we propose that the Las-quorum-sensing regulon might no longer be beneficial under many of the culturing conditions used in the laboratory, especially when bacteria are consistently grown at high cell densities. Mutations in *lasR* would thus reflect the first step in the degradation of this system. Alternatively, it is conceivable that quorum sensing remains beneficial, but that mutations in *lasR* represent the first step in the rewiring of the QS network in order to customize it to the novel conditions experienced in infections and laboratory cultures [67]. Finally, the invasion of *lasR*-mutants could be the result of cheating, where these signal blind mutants still contribute to signal production, but no longer respond to it and thus refrain from producing the QS-controlled public goods [47,68]. We have argued above that, although *lasR* mutants were favoured in all our treatments, the presence of the host might change the selection pressure and underlying reason for why these mutants are selected for. More generally, our observations of high strain diversification during experimental evolution, and the co-existence of multiple different phenotypes and genotypes within each replicate, are reminiscent of patterns found in chronic *P. aeruginosa* infections in cystic fibrosis lungs [19,61,62,69–72]. While this diversity might be transient in some cases, it highlights that an initially clonal infection can give rise to a diverse community, with multiple strains competing with each other within the host, as it was observed in CF lung communities [73,74]. Despite these striking similarities, we need to be careful when extrapolating from a nematode gut to a human lung environment. Clearly, more studies in other host organisms are required to identify common evolutionary patterns in infections. Moreover, analysis of intermediate time points and additional virulence factors could further deepen our understanding of temporal evolutionary patterns and virulence traits under selection.

In conclusion, our study demonstrates that there is rapid and parallel virulence evolution in populations of the opportunist *P. aeruginosa*, and that secreted virulence factors are the main target of selection. While low spatial structure of the environment generally selected for lower virulence regardless of whether hosts were present or not, the virulence traits under selection and the strength of selection were host dependent. This greatly contributes to our knowledge on how bacterial opportunistic pathogens adapt to the variable environments they occupy, and how this affects their virulence [26,27]. Our work also highlights that linking virulence evolution to selection inside and outside of the host is key to predict evolutionary trajectories in opportunistic pathogens. Such insights might offer simple approaches of how to manage infections in these clinically highly important pathogens [69,75–77], for example through the disruption of spatial structure in chronic infections, which could, according to our findings, steer pathogen evolution towards lower virulence.

## DATA AVAILABILITY

All sequencing data generated for this study are available from the European Nucleotide Archive (accession number PRJEB23190). All other raw datasets have been deposited in the Figshare repository (doi:).

## SUPPLEMENTARY INFORMATION

Supplementary information is available at ISME’s website.

## ACKNOWLEDGEMENTS

We thank Swati Parekh for assistance in genomic analysis, Isa Moreno for help with the nematode killing assays, Ruben Dezeure for statistical advice, and Stu West and Sam Brown for comments on the manuscript.

